# Uncovering the Elusive Structures and Mechanisms of Prevalent Antidepressants

**DOI:** 10.1101/2024.01.04.574265

**Authors:** Jieye Lin, Guanhong Bu, Johan Unge, Tamir Gonen

**Author notes:** Corresponding Author T.G.

## Abstract

Most treatments to alleviate major depression work by either inhibiting human monoamine transporters, vital for the reuptake of monoamine neurotransmitters, or by inhibiting monoamine oxidases, which are vital for their degradation. The analysis of the experimental 3D structures of those antidepressants in their drug formulation state is key to precision drug design and development. In this study, we apply microcrystal electron diffraction (MicroED) to reveal the atomic 3D structures for the first time of five of the most prevalent antidepressants (reboxetine, pipofezine, ansofaxine, phenelzine, bifemelane) directly from the commercially available powder of the active ingredients. Their modes of binding are investigated by molecular docking, revealing the essential contacts and conformational changes into the biologically active state. This study underscores the combined use of MicroED and molecular docking to uncover elusive drug structures and mechanisms to aid in further drug development pipelines.

## Introduction

Major depression is a complicated mood disorder caused by biological, psychological and social factors.^1,2^ Based on the monoamine hypothesis,^3,4^ current therapy manipulates the availability and activity of three monoamine neurotransmitters (norepinephrine, NE; serotonin, 5-HT; and dopamine, DT) in synapses. Normal neuron-neuron communication encompasses the release and binding of monoamine neurotransmitters from the presynaptic neuron to receptors in the postsynaptic neuron. The monoamine neurotransmitters that remain in the synaptic cleft can be subject to reuptake through monoamine transporters (human norepinephrine transporter, hNET; human serotonin transporter, hSERT; human dopamine transporter, hDAT)^5^. The excess is degraded by monoamine oxidase A/B (MAO-A/B)^6^ in the presynaptic neuron (Fig. 1a). Antidepressants that inhibit monoamine transporters or oxidases involved in these processes help increase the bioavailability of monoamine neurotransmitters (Fig. 1b). For example, reboxetine is a norepinephrine reuptake inhibitor (NRI) inhibiting the function of hNET.^7,8^ Pipofezine is a tricyclic antidepressant (TCA) functioning on hSERT^9,10^ while ansofaxine interacts with hSERT, hNET, hDAT and is a serotonin–norepinephrine–dopamine reuptake inhibitor (SNDRI).^11^ Monoamine oxidase inhibitors (MAOIs) like phenelzine^12,13^ inhibit the MAO-A/B irreversibly, while bifemelane^14,15^ interact in a reversible mode.

**Fig. 1.**
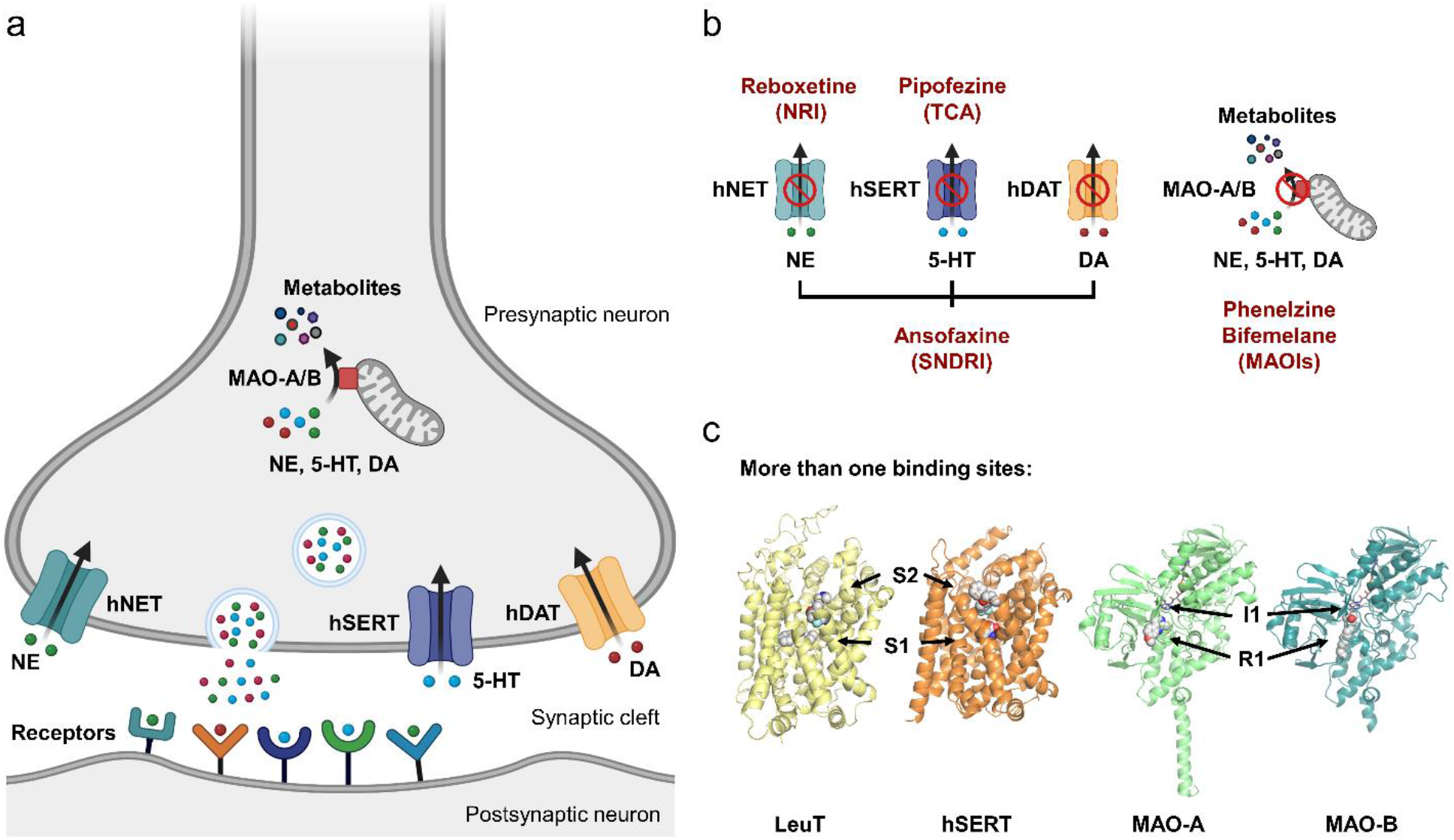
(a) Reuptake and degradation of NE, 5-HT, and DA in neuron-neuron communication. (b) hNET, hSERT, hDAT and MAO-A/B may be inhibited by five antidepressants. (c) The PDB structures reveal more than one binding sites in hNET, hSERT, and hDAT (central: S1; allosteric: S2), and in MAO-A/B (irreversible: I1; reversible: R1) for antidepressants. Created with BioRender.com
.

The experimental 3D structure of antidepressants reveals the energetically favorable conformations, which determines the precise properties of the formulations used for drug distribution, and which are fundamental for modeling the energetic and conformational changes needed for binding to the protein pocket. In structure-based drug design these insights are essential for precision drug design and development. To date, most of the crystal structures of antidepressants have been solved by single-crystal X-ray diffraction (SC-XRD), like desvenlafaxine,^16^ viloxazine,^17^. A fundamental step needed for SC-XRD is to obtain large enough crystals (>5 μm) to have adequate diffraction.^18^ This is however sometimes unattainable due to the imperfect crystals. Techniques like powder X-ray diffraction (PXRD)^19^ and solid-state nuclear magnetic resonance (ssNMR)^20,21^ may be challenging due to peak overlapping and broadening, and extensive calculations may be required. Therefore, several antidepressant compounds remain without a known crystal structure for several years. The new cryogenic electron microscopy (Cryo-EM) technique microcrystal electron diffraction (MicroED) serves as a complementary route for experimental crystal structure determination,^22,23^ which enables the direct analysis of microcrystals from a seemingly amorphous powder, where the crystal size needed is merely a billionth of the size needed for SC-XRD.^24^ Since the advent of this technique already delivered many novel drug structures, like antihistamines^25,26^ and macrocyclic drugs^27,28^.

Despite the variety of antidepressants, the knowledge of their interactions with target proteins remains limited. The biggest hurdle for the determinations of the complex structures is the purification and stabilization of their target receptors: membrane proteins like hNET and hDAT.^29^ As a result, model proteins such as the drosophila dopamine transporter (dDAT) and the bacterial leucine transporter (LeuT) were used as homology models for the human proteins. Their binding complexes with antidepressants have been reported, such as for reboxetine/dDAT^30^ and sertraline/LeuT.^31^ Although only the structures have 56% and 25% sequence identity to the human sequences, respectively, they interestingly indicate more than one binding site for the antidepressants on their respective target (Fig. 1c). However, the human counterparts vary in sequence affecting the exact recapturing the binding mode in humans, possible residue changes in active sites remain unknown and may effect drug binding. Although the complex structures of hSERT and selective serotonin reuptake inhibitor (SSRI) like citalopram or paroxetine have been published (PDB [PDB, https://www.rcsb.org/] entries: 5I73, 5I6X),^32^ the structures of the complexes of hSERT with pipofezine (TCA) or ansofaxine (SNDRI) remain undetermined, as is also the case for the complexes of MAO-A/B to phenelzine and bifemelane (MAOIs).

In this study, we applied MicroED to determine the atomic structures of five prevalent commercial antidepressants, reboxetine (Europe, 1997),^8^ pipofezine (Russia, 1960),^10^ ansofaxine (China, 2022),^11^ phenelzine (United States, 1961)^13^ and bifemelane (Japan, 1980)^15^ which mostly have remained undetermined for decades although widely prescribed worldwide (Fig. 1b). To uncover their binding mechanism in human targets, molecular docking was applied using the ligand structures determined by MicroED and the receptor structures from either the Protein Data Bank or from AlphaFold predictions.^35^ The docked target-bound structures were analyzed to show key interactions with the residues in the active sites, and their conformational changes from the drug formulation state.

## Results and Discussion

### Antidepressant structures in their drug formulation state

For each antidepressant, just a few nanograms of powder were used in MicroED sample preparation without recrystallization (See Methods for more details).^24^ A Thermo-Fisher Talos Arctica transmission electron microscope (TEM) (200 keV) equipped with CetaD CMOS camera (4096 × 4096 pixels) and EPU-D software (Thermo-Fisher) was used for MicroED data collection. The microcrystal screening was conducted in imaging mode (SA 5300x), once suitable nanocrystals were identified, their eucentric heights were adjusted to keep the sample within the beam during rotation. The crystals of five antidepressants all appeared as thin plates or sheets (Fig. 2a-e) which are not ideal for SC-XRD, possibly explaining why their x-ray crystal structures have remained elusive for decades. Continuous rotation MicroED data collection^23^ was conducted in the diffraction mode and the parallel electron beam settings using an electron dose rate (exposure) of 0.01 e^-^/(Å^2^·s). MicroED data were recorded as movies at 0.5 s exposure per frame as the microcrystal was continuously rotated at 2° per second across a 120° angular wedge (from -60° to +60°), thus the total dose for each dataset was ∼0.6 e^-^/Å^2^. The MicroED movies were converted to images using mrc2smv.^36^ Diffraction data was indexed, integrated, scaled, and merged in XDS^37,38^ to generate the reflection files which could be *ab initio* solved by SHELXT^39^/SHELXD^40^ and refined by SHELXL^41^ (Fig. 2f and Fig. 3, Supplementary Table 1).

**Fig. 2.**
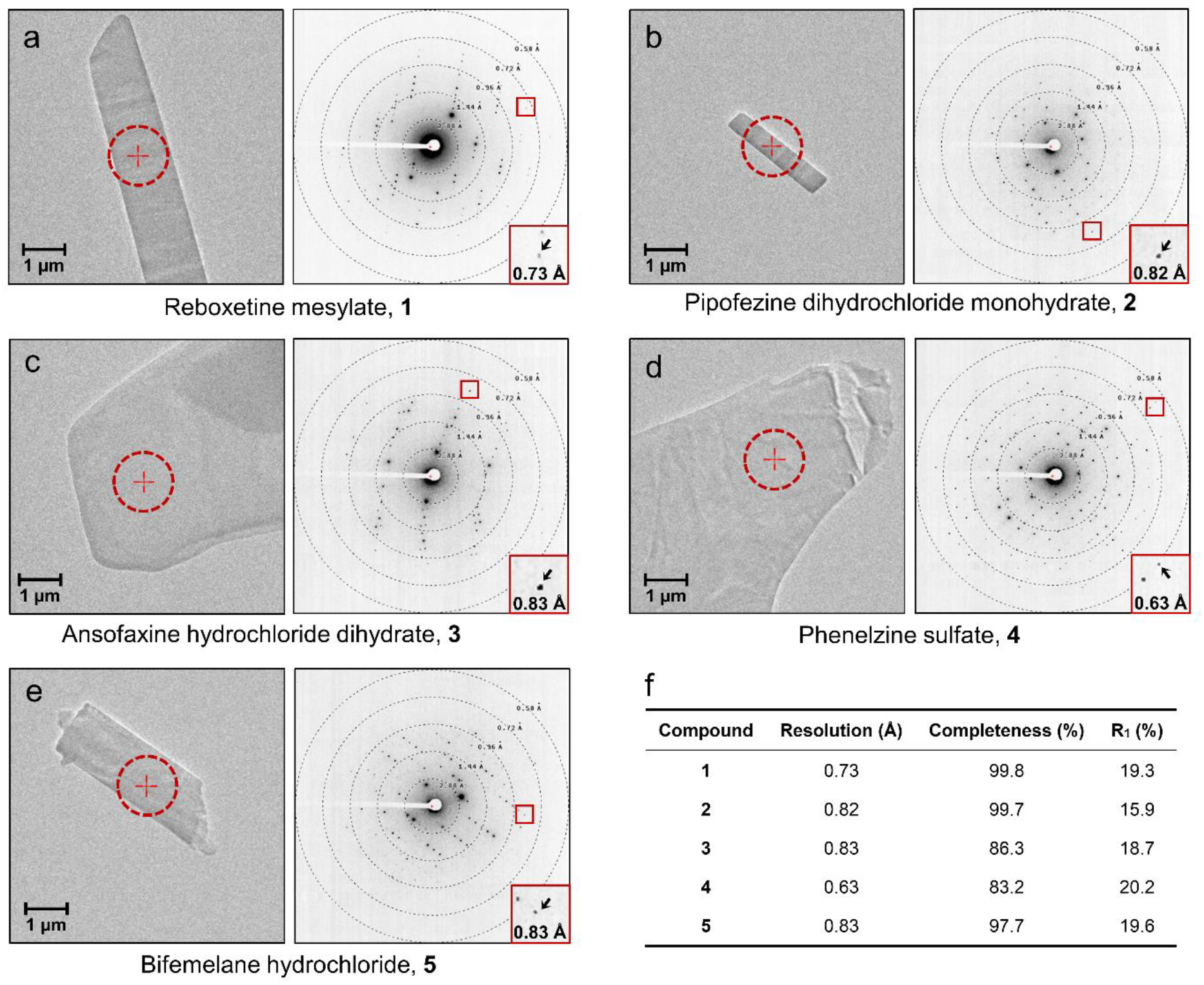
(a-e) Images of the crystals together with typical MicroED patterns of **1**-**5**. (f) MicroED data statistics of **1**-**5**, see more details in Supplementary Table 1.

**Fig. 3.**
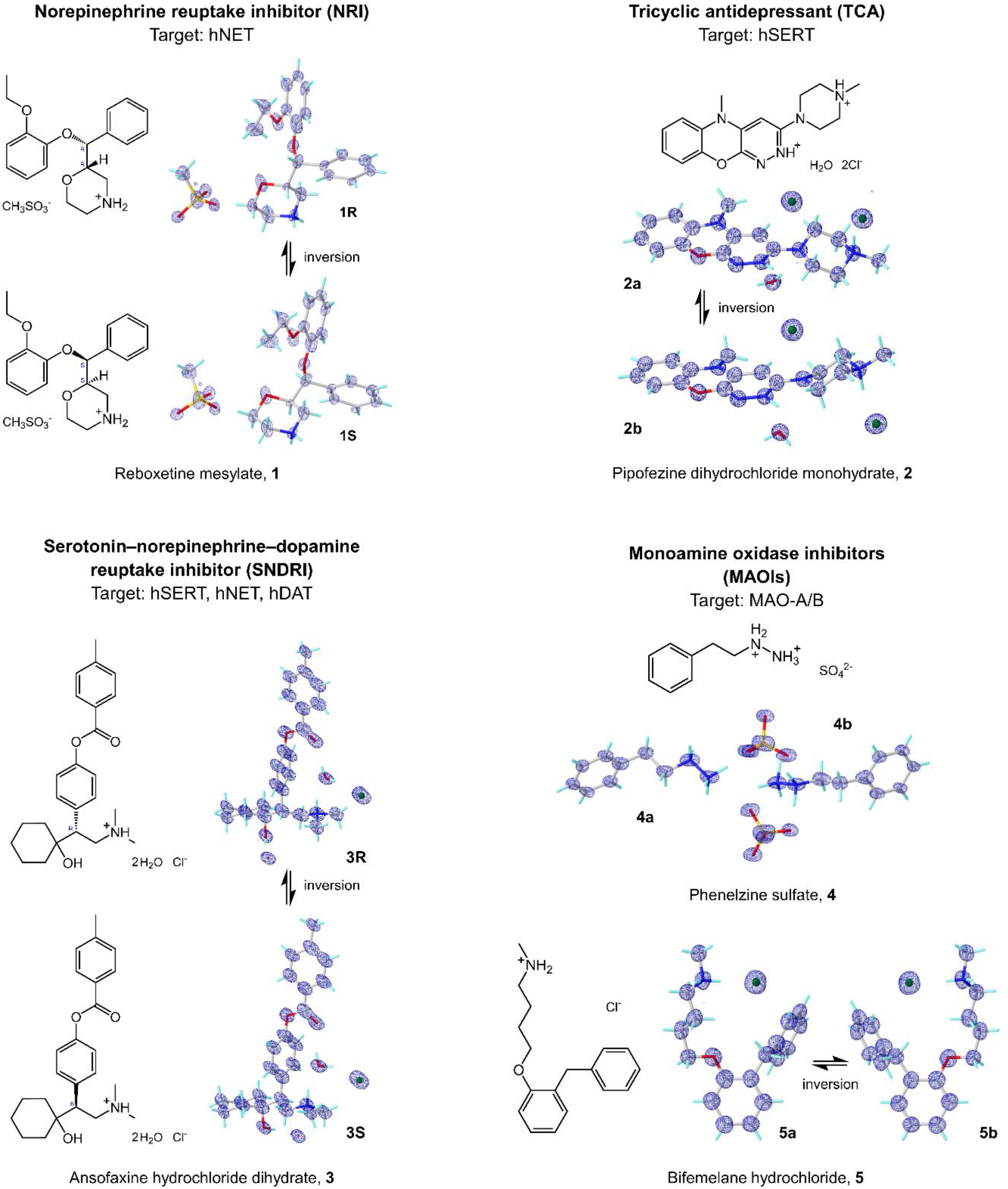
MicroED structures and 2F_o_-F_c_ density maps (blue mesh) of **1**-**5. 1R**/**1S, 2a**/**2b, 3R**/**3S** and **5a**/**5b** are related by an inversion symmetry in the unit cell, **4a**/**4b** are two conformers observed in one asymmetric unit. Atom color: C, gray; N, blue; O, red; S, yellow; Cl, green; H, light green.

The atomic MicroED structures of five antidepressants: reboxetine mesylate **1**, pipofezine dihydrochloride monohydrate **2**, ansofaxine hydrochloride dihydrate **3**, phenelzine sulfate **4**, and bifemelane hydrochloride **5**, have been solved for the first time (Fig. 3). These elusive crystal structures offered detailed structural information about their drug formulation state, including conformers, anhydrous/hydrate forms, and salts. These may serve as a starting point for understanding the conformational changes upon binding to the target pockets in their biologically active states.

1. **Reboxetine mesylate 1**. The racemic MicroED structure **1** was solved in a centrosymmetric monoclinic P 2_1_/c space group at the resolution of 0.73 Å (Fig. 2a and Fig. 3), with the unit cell parameters of **a** = 20.00 Å, **b** = 5.49 Å, **c** = 19.06 Å, **α** = 90.000°, **β** = 107.338°, **γ** = 90.000°. Two enantiomers, namely **1R** and **1S**, stack their hydrophobic and hydrophilic parts in layers along the *c*-axis so that the layers are parallel with the *a*- and *c*-plane. Within these layers, molecules adopt a head-to-tail orientation along the *a*-axis, linked by mesylate anions (Supplementary Fig. 1). The crystal packing is mainly a result of several hydrogen bonds and dipole-dipole interactions along *b*- and c-axes, for instance the N2/N2′-H···O6 (3.30 Å) bond to the sulfonyl group (atom O6) and the dipole-dipole interactions involving the CH atoms in reboxetine and O atoms in mesylate. The elongation along *a*-axis is stabilized by T-shaped/parallel-displaced pi-stacking interactions ranging from 4.81 to 5.50 Å. In total six torsion angles are freely rotatable, which dramatically impact the whole structure (Supplementary Fig. 2).
2. **Pipofezine dihydrochloride monohydrate 2**. The MicroED structure **2** was solved in a monoclinic P 2_1_/c space group at the resolution of 0.82 Å (Fig. 2b and Fig. 3), with the unit cell parameters of **a** = 6.88 Å, **b** = 15.61 Å, **c** = 15.93 Å, **α** = 90.000°, **β** = 97.222°, **γ** = 90.000°. Two conforms, namely **2a** and **2b** were identified in the unit cell. Each can be transformed by inversion symmetry or 180° rotation of C11-N4/C11′-N4′ bond. The crystal packing is formed mainly by hydrogen bonds and ion-dipole interactions between **2a/2b** and chloride anions along *b*- and *c*-axes, *i*.*e*. hydrogen bonds N5/N5′-H···Cl1 (3.01 Å) and N3/N3′-H···O2 (2.67 Å); ion-dipole interactions between CH atoms and chloride anions (Supplementary Fig. 3). The water molecules serve as hydrogen bond donors to Cl1 or Cl2 anions that bridge **2a** and **2b** molecules together (Supplementary Fig. 3). The packing along the *a*-axis is facilitated by strong parallel-displaced pi-stacking interactions between the phenyl and pyridazine rings in **2a** and **2b** (3.65 Å). In **2**, bond angles are mostly fixed, with only one freely rotating bond (C10-C11-N4-C15 and C10′-C11′- N4′-C15′, measured at ±178.60° in **2a** and **2b**), generating a co-planar arrangement of piperazine ring and tricyclic moiety (Supplementary Fig. 4).
3. **Ansofaxine hydrochloride dihydrate 3**. The racemic MicroED structure **3** was solved in a centrosymmetric monoclinic P 2_1_/c space group at the resolution of 0.83 Å (Fig. 2c and Fig. 3), with the unit cell parameters of **a** = 14.80 Å, **b** = 10.27 Å, **c** = 16.04 Å, **α** = 90.000°, **β** = 95.309°, **γ** = 90.000°. Two enantiomers, namely **3R** and **3S**, pack as *zig-zag* shaped layers along the *c*-axis. Within each layer, molecules orient head-to-tail along the *a*-axis, interconnected by two water molecules and a chloride anion (Supplementary Fig. 5). Each water molecule involves in two hydrogen bonds as the acceptor and another two hydrogen bonds as the donor, the Cl^-^ anion contains four lone pair electrons that can be the acceptor for four hydrogen bonds, that leads to a dense hydrogen bonding network. The packing along the *b*- and *c*-axes involves extensive hydrogen bonding interactions, *i*.*e*., N1/N1′-H···O5 (2.80 Å), O5-H···O1/O1′ (2.85 Å) O4-H···Cl1 (3.02 Å) along the *b*-axis; O3/O3′-H···O4 (2.73 Å) and O4-H···O5 (2.81 Å) along the *c*-axis. Although the O5-H···Cl1 (3.01 Å) orients along the *a*-axis, it cannot further elongate the crystal packing, the latter is largely achieved by medium or weak T-shaped pi-stacking interactions between **3R** and **3S** ranging from 5.25 to 5.88 Å. In **3**, bond lengths and angles are rather fixed, with six torsion angles exhibit a high degree of rotational freedom (Supplementary Fig. 6).
4. **Phenelzine sulfate 4**. The MicroED structure **4** was solved in a centrosymmetric monoclinic P 2_1_/c space group at the resolution of 0.63 Å (Fig. 2d and Fig. 3), with the unit cell parameters of **a** = 20.36 Å, **b** = 5.46 Å, **c** = 20.30 Å, **α** = 90.000°, **β** = 111.209°, **γ** = 90.000°. Two conformers, namely **4a** and **4b**, were found in the asymmetric unit. As the molecule does not possess a chiral center, the inversion symmetry results in slight differences in the torsion angles (maximum 14° difference, see Supplementary Fig. 8). The positively charged nitrogen containing moieties of **4a** and **4b** forms a hydrophilic network together with the sulfate anions. This network distributes as layers parallel to the b-c plane separated by the hydrophobic layers containing the aromatic and aliphatic part of the molecules. The molecules in the layers adopt a head-to-tail orientation similar to lipid layers. The hydrophilic network is constructed by several hydrogen bonds between **4a/4b** and sulfate anions, such as N1-H···O5/O8 (2.73 Å/2.66 Å), N2-H··· O1/O4/O6 (2.73 Å/2.73 Å/2.67 Å), N1′-H···O3/O7 (2.59 Å/2.65 Å), N2′-H···O2/O4/O5 (2.87 Å/2.65 Å/2.75 Å) (Supplementary Fig. 7). The T-shaped/parallel-displaced pi-stacking interactions ranging from 5.02 to 5.93 Å facilitate the packing along the *a*-axis. Bond angles are mostly fixed, with three freely rotatable torsion angles corresponding to similar values in two conformers. In the biological state, only the phenylethyl part will covalently bind to the flavin adenine dinucleotide (FAD) cofactor in MAO-A/B, where the C4-C7/C4′-C7′ is presumed to have 44° to 69° rotations, see details below (Fig. 5a, b).
5. **Bifemelane hydrochloride 5**. The MicroED structure **5** was solved in a centrosymmetric orthorhombic P bca space group at the resolution of 0.83 Å (Fig. 2e and Fig. 3), with the unit cell parameters of **a** = 12.87 Å, **b** = 7.31 Å, **c** = 35.10 Å, **α** = 90.000°, **β** = 90.000°, **γ** = 90.000°. Two conformers, namely **5a** and **5b** forms hydrophobic and hydrophilic layers parallell with the *a*- and *b*-plane. A double-helical extension was observed along the *c*-axis. Complex hydrogen bonds between **5a/5b** and chloride anions dominate the packing along the *a*- and *b*-axes, for example, the N1-H1N···Cl1 and N1′-H1N′···Cl1 (3.04 Å) along the *a*-axis; the bifurcated N1-H2N···Cl1 and N1′-H2N′···Cl1 (3.10/3.58 Å) are along the *a*- and *b*-axes. However, the packing along the *c*-axis is facilitated by the T-shaped pi-stacking interactions between two phenyl rings (5.09 to 5.98 Å). In the bifemelane structure, as many as eight torsion angles are freely rotatable which can significantly change the overall structure (Supplementary Fig. 10). The high number of freely rotatable bonds enhance the conformational flexibility, facilitating the binding within the protein pockets.

### Binding mechanisms of 1-5 predicted by molecular docking

In the docking setup, the MicroED ligand structures of **1**-**3** and **5** were used. Since only the phenethyl part of **4** covalently binds to FAD cofactor in MAO-A/B, a substitute structure **4\*** was extracted from PDB entry 2VRM.^44^ Three X-ray structures: hSERT (PDB entry: 5I73),^32^ MAO-A (PDB entry: 2Z5X),^33^ MAO-B (PDB entry: 1OJ9),^34^ along with two AlphaFold structures^35^: hNET (PDB entry: AF_AFP23975F1) and hDAT (PDB entry: AF_AFQ01959F1) were utilized. The AlphaFold structures of hNET and hDAT were calculated from human SLC6A2 and SLC6A3 gene sequences respectively with high scores. The binding sites in hNET, hSERT, hDAT and MAO-A/B have been reported in literature describing the complex structures (Fig. 1c; Supplementary Fig 11, Supplementary Table 2).^5^ During the molecular docking, the ligands were set to be flexible while keeping the receptor rigid. The docked complex with the lowest energy was examined (see details in Methods).

#### (1) Inhibition of the reuptake process of NE, 5-HT and DA by 1-3

##### 1-hNET complexes

**1R** and **1S** were docked at hNET in the central (S1) and allosteric (S2) sites respectively (Supplementary Fig. 12a-b),^30,31^ their binding interactions are similar but differ in certain cyclic moieties. For example, within the S1 site, **1R** is primarily bound by two hydrogen bonds between the morpholine ring and Phe317, Ser419 residues. Another pi-stacking (Tyr152) and nine hydrophobic interactions stabilize the conformation of phenyl and 2-ethoxyphenoxy rings (Fig. 4a). The binding interactions observed in **1R**/dDAT complex (PDB: 4XNX)^30^ exhibit weaker hydrogen bonds and fewer hydrophobic interactions. **1S** in the S1 site is bound by two hydrogen bonds (Asp75, Tyr152), one pi-stacking (Tyr152) and nine hydrophobic interactions (Fig. 4b). The conformation changes of **1R** and **1S** at S1 site were predominantly driven by the rotations of C18-O3, C5-C9 and C5′-C9′ bonds (Supplementary Fig. 2).

**Fig. 4.**
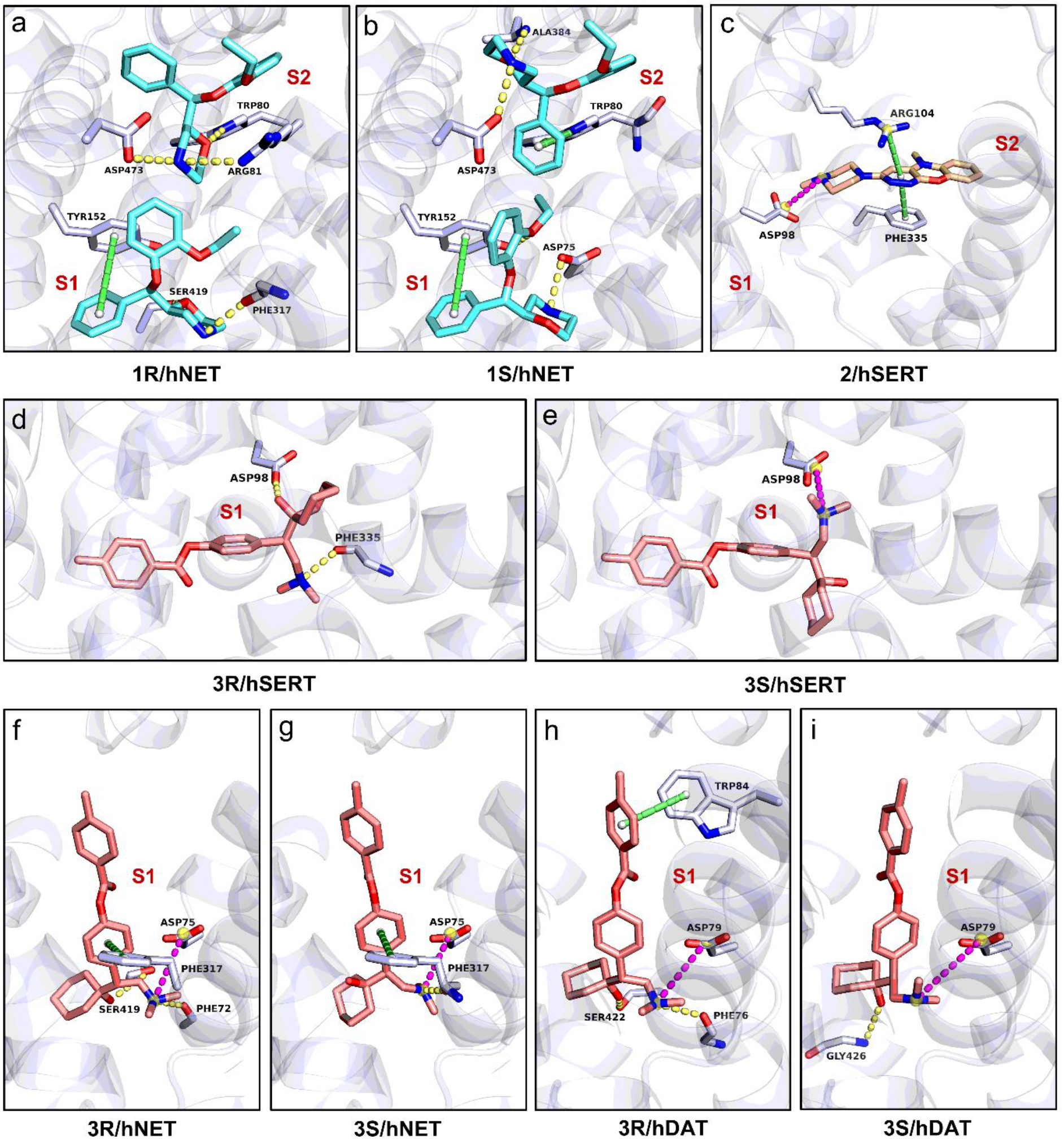
The protein-drug interaction diagram of complex between (a) **1R** and hNET; (b) **1S** and hNET; (c) **2** and hSERT; (d) **3R** and hSERT; (e) **3S** and hSERT; (f) **3R** and hNET; (g) **3S** and hNET; (h) **3R** and hDAT; (i) **3S** and hDAT. Hydrogen bonding interactions were colored by the dashed line in paleyellow, pi-stacking interactions were colored by the dashed line in lime, and salt bridges were colored by the dashed line in magenta, hydrophobic interactions were omitted for clarification. The binding sites were marked, see the overall view in Supplementary Fig. 12.

**Fig. 5.**
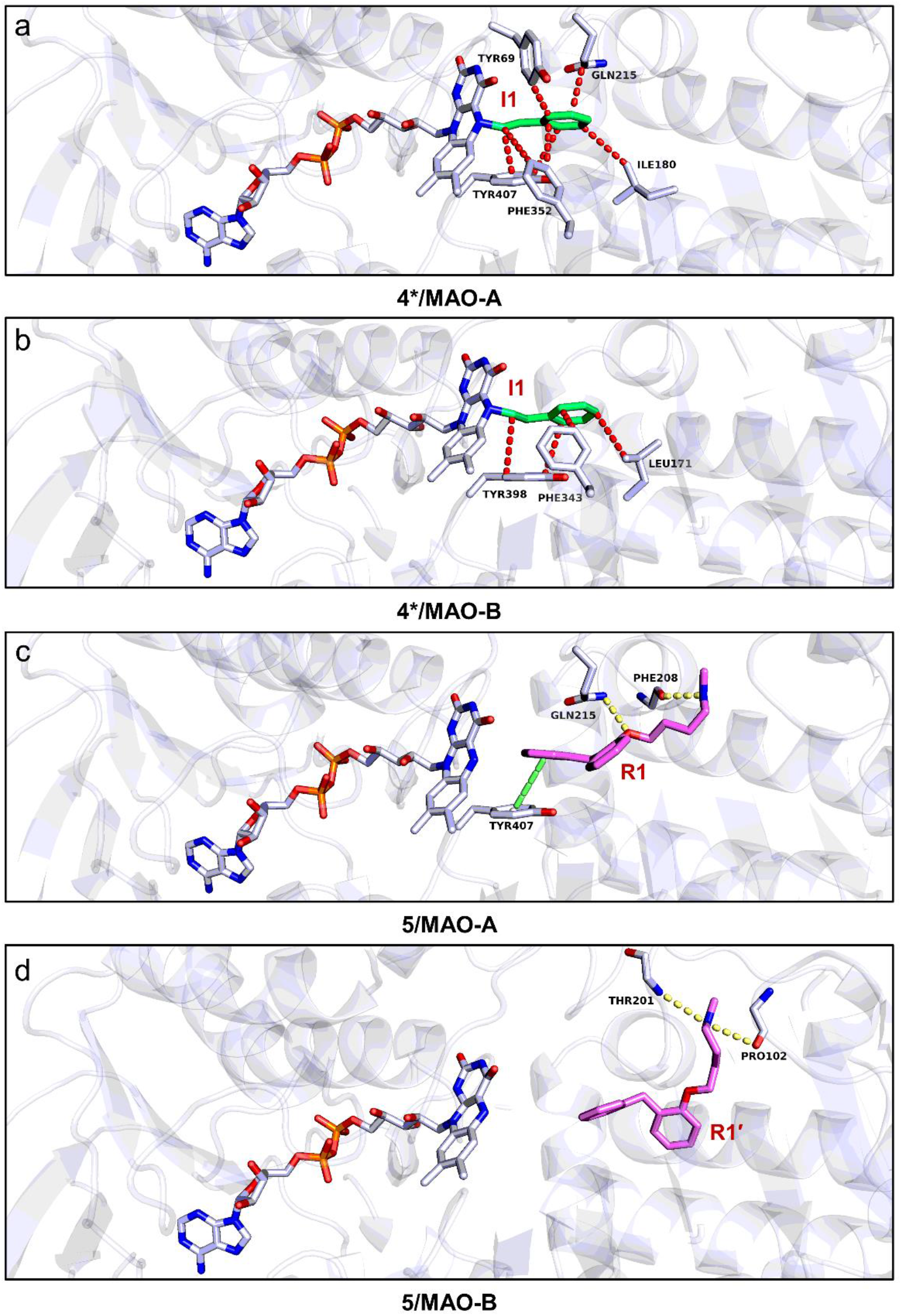
The protein-drug interaction diagram of complex between (a) **4\*** and MAO-A; (b) **4\*** and MAO-B; (c) **5** and MAO-A; (d) **5** and MAO-B. Hydrogen bonding interactions were colored by the dashed line in paleyellow, pi-stacking interactions were colored by the dashed line in lime. Hydrophobic interactions were colored by the dashed line in red for **4\*** but omitted for **5**. The binding sites were marked, see the overall view in Supplementary Fig. 13.

The binding interactions in the S2 site seems comparably “tight” as the one in the S1 site. For instance, **1R** is bound by three hydrogen bonds between the morpholine ring and Trp80, Arg81, Asp473 residues, and five hydrophobic interactions for the rest of structure (Fig. 4a); whereas **1S** is bound by two hydrogen bonds (Ala384 and Asp473), two pi-stacking (Trp80), and five hydrophobic interactions (Fig. 4b). The conformation changes of **1R** and **1S** at S2 site were mainly attributed to the rotations of C18-O3, C12-O2, C13′-O3′ and C9′-C5′ bonds (Supplementary Fig. 2). The S1 and S2 sites have a comparable amount of interactions with the molecule indicating activity at both suggested sites, which fits well with the reported view of S1 being the central site and S2 an allosteric site. When the central site of **1R**/**1S** is occupied, the hNET is maintained in the outward-open conformation and obstruct the reuptake channel of NE.^5^

##### 2-hSERT complexes

**2** was tested at the central (S1) and allosteric (S2) sites in hSERT (Supplementary Fig. 12c),^32^ however the final docking center was found near S2 site due to the weak binding observed in S1 site (*i*.*e*. only the hydrophobic interactions). A salt bridge between Asp98 and piperazine ring, together with one pi-stacking (Phe335), one pi-cation interaction (Arg104), three hydrophobic interactions (Phe335, Phe556) to the tricyclic moiety stabilized the binding complex of **2**/hERT (Fig. 4c). The structures of **2** in its drug-formulation state and biologically active state are very similar, with only 3-6° rotation C11′-N4′ bond (Supplementary Fig. 4), and maintain the piperazine ring and tricyclic moiety in a nearly co-planar geometry (C10-C11-N4-C15≈180°) for both states. The minimum conformational changes ensure small entropy differences upon binding which is beneficial for the binding of **2** to the receptor.

##### 3-hSERT, hNET, hDAT complexes

**3R**/**3S** demonstrated inhibitions against hSERT, hNET and hDAT as a triple reuptake inhibitor.^11^ Due to their larger molecular occupancy, only the central (S1) site was docked (Supplementary Fig. 12d-i). The binding interactions between **3R/3S** and hSERT, hNET, hDAT differ because of the structure and sequence variations in their binding pockets.

In **3R/3S**-hSERT complexes, **3R** is involved in two hydrogen bonds between the dimethyl amine group and Phe335 residue, and between the 1-hydroxycyclohexyl ring and Asp98 residue (Fig. 4d). **3S** also adopts a salt bridge between the dimethyl amine group and Asp98 residue (Fig. 4e). In addition, the binding of **3R/3S** is stabilized by seven and eleven hydrophobic interactions with the target respectively. The above conformation can be reached by a simple rotation of C12′-C15′ bond in **3S**, while four bond rotations (C5-C8, C9-O2, C12-C15, C22-N11) were required in **3R** during the transition to its biologically active state (Supplementary Fig. 6).

In **3R/3S**-hNET complexes, **3R** is bound by one salt bridge (Asp75), two hydrogen bonds between the dimethyl amine group and Phe72, and the 1-hydroxycyclohexyl ring and Ser419, respectively. The additional pi-stacking (Phe317) and nine hydrophobic interactions stabilized the rest of the structure (Fig. 4f). The similar binding was found for **3S**, where the dimethyl amine group is bound by one salt bridge (Asp75) and one hydrogen bond (Phe317), with the rest of the molecule anchored by one pi-stacking (Phe317) and eight hydrophobic interactions (Fig. 4g). To transition from drug-formulation state to biologically active state, **3R** requires rotations at C5-C8, C12-C15, and C22-N1 bonds, while **3S** requires changes at C5′-C8′, C12′-C15′, and C15′-C16′ bonds (Supplementary Fig. 6).

In **3R/3S**-hDAT complexes, **3R** engages in a rigid salt bridge (Asp79) and two hydrogen bonds (Phe76, Ser422) as similar to hNET; the rest of molecule is anchored by one pi-stacking (Trp84) and up to eleven hydrophobic interactions (Fig. 4h); Similarly, **3S** is bonding with a salt bridge between the dimethyl amine group and Asp79 residue, one hydrogen bond between the 1-hydroxycyclohexyl ring and Gly426 residue, and twelve hydrophobic interactions (Fig. 4i). The conformational shifts from drug formulation state to biologically active state needs two major bond rotations, like C12-C15 and C22-N1 in **3R** and C9′-O2’ and C22′-N1′ in **3S** (Supplementary Fig. 6).

#### (2) Inhibition of the degradation process of NE, 5-HT and DA by 4-5

##### 4*-MAO-A/B complexes

Under the catalysis of MAO-A/B, the N-H bond in **4** will be cleaved to result a diazene intermediate.^44^ The diazene was oxidized to produce an arylalkyl radical, which reacted rapidly with N(5) atom of flavin group (I1 site) and formed an adduct in FAD as **4\*** (Supplementary Fig 13a,b).^12,44^ The enzyme function can be inhibited because of the inactivation of FAD. The crystal structure of **4\***/MAO-B was reported (PDB entry: 2VRM),^44^ where the major rotation in phenethyl part was found in C4-C7/C4′-C7′ bond, from 107-109° to 176°, leading the co-planar orientation of C7-C8 bond and phenyl ring (Supplementary Fig. 8). Such conformation was maintained by three hydrophobic interactions between the phenethyl group and Leu171, Phe343, Tyr398 residues (Fig. 5b). Since the position of FAD in MAO-A and MAO-B are highly conserved, the **4\*** was extracted and docked in MAO-A. The resulting model is very similar to **4*/**MAO-B, showing the co-planar of C7-C8 bond and phenyl ring (Supplementary Fig. 8), and five hydrophobic interactions between the phenethyl group and Tyr69, Ile180, Gln215, Phe352, Tyr407 residues (Fig. 5a).

##### 5-MAO-A/B complexes

The reversible inhibitor **5** was proven to bind with a higher inhibition of MAO-A (K_i_=4.2 μM) than MAO-B (K_i_=46.0 μM).^14^ The binding site of **5**/MAO-A was docked in the active site (R1 site, Supplementary Fig. 13c),^33^ where **5** was bound by two hydrogen bonds between the *N*-methylbutylamine group and the Phe208 residue, and between the phenoxyl ring and Gln215 residue, respectively. One pi-stacking (Tyr407) and up to ten hydrophobic interactions restrain the rest of molecules (Fig 5c). The above conformational shifts from the drug formulation state were completed by the rotation of C14-O1, C14-C15 and C16-C17, making the molecule be stretched in the large voids of MAO-A (Supplementary Fig. 10).

In contrary to MAO-A, the active site in MAO-B is surrounded by several hydrophobic residues, and more restricted space at the entrance impede the direct entry of large molecules (Supplementary Fig. 14). As a result, **5** was trapped at the R1′ site (Supplementary Fig. 13d),^14,33^ which has ∼12.3 Å distance to the N(5) atom of FAD. Such distance in MAO-A is around 8.8 Å (Supplementary Fig. 14), which possibly demonstrates the higher affinity of **5** to MAO-A rather than MAO-B. In the R1′ site, **5** was bound by two hydrogen bonds (Pro102, Thr201) and ten hydrophobic interactions (Fig 5d), keeping it in a more folded conformation. The above conformation was completed by the rotation of C14-O1, C14-C15, C16-C17, C17-N1 in the *N*-methylbutylamine group (Supplementary Fig. 10).

## Conclusions

Characterizing the 3D structures of antidepressants from their drug formulation state and the biologically active state are both essential for precision drug design and development. Conventionally, SC-XRD has been the most frequently used technique for drug structure elucidation, which however requires larger well-ordered crystals (>5 μm). Other techniques like PXRD and solid-state NMR have been important for structure determinations of smaller molecules, but larger targets are associated with problems in peak broadening or overlapping, and require extensive computational simulation to deliver an atomic model.^19-21^ Several important targets such as the five antidepressants analyzed in this study do not easily yield large enough crystals for SXR-XD or PXRD so their 3D structures remained elusive for decades. The emerging Cryo-EM technique MicroED^22,23^ serves as a complementary route for experimental structure determination directly from powder-like microcrystals^24-28^, which are the prevalent form of formulated drugs or active pharmaceutical ingredients. Thus, it may bypass the time and resources spent in efforts to produce large crystals for SC-XRD.

Analysis of the crystal packing in the five structures **1**-**5** revealed a difference in packing interactions along three dimensions in any of the crystals. For example, in **3**, the extensive hydrogen bonding interactions (2.73 to 3.02 Å) extend the packing in *b*- and *c*-axes, but T-shaped pi-stacking interactions (5.25 to 5.88 Å) mainly correspond for packing elongation along *a*-axis. These differences in packing in different direction in the crystal potentially translate to differences in crystal growth along the different directions and hence may result in crystals being plates or sheets due to a slower growing expansion of the crystals in the third dimension. Such crystal shapes, with a large area but a thickness far below one micron are unsuitable for SC-XRD but ideal for MicroED.

Although it is important to obtain protein-ligand complex structures for **1**-**5** experimentally, the purification and crystallization of those membrane proteins are extremely difficult and they are too small for single particle cryoEM analyses without bulking up their size with deactivating antibodies for example. Instead, molecular docking was used here as an alternative technique to predict the interactions between **1**-**5** and their target proteins directly using the MicroED structures of **1**-**5**. This approach revealed unknown properties in both drug and proteins, for example, the binding orientations of **3R/3S** in hSERT, hNET, hDAT were determined by a salt bridge between Asp residue and dimethyl amine group (Fig. 4d-i); The structural differences found in MAO-A/B led to different selectivity with **5**. The conformational changes of **1**-**5** from drug formulation state to the biologically active state were also elucidated, pinpointing to the possible geometric compromises caused by the rotation of dynamic groups within the drugs. This study underscored the combined use of MicroED and molecular docking in uncovering elusive antidepressant drug structures and mechanisms which are crucial for the future drug design and development.

## Acknowledgements

The authors thank Michael W. Martynowycz for support and discussions. This study was funded in part by the National Institutes of Health P41GM136508. Portions of this research or manuscript completion were developed with funding from the Department of Defense grants MCDC-2202-002. Effort sponsored by the U.S. Government under Other Transaction number W15QKN-16-9-1002 between the MCDC, and the Government. The US Government is authorized to reproduce and distribute reprints for Governmental purposes, notwithstanding any copyright notation thereon. The views and conclusions contained herein are those of the authors and should not be interpreted as necessarily representing the official policies or endorsements, either expressed or implied, of the U.S. Government. The PAH shall flowdown these requirements to its subawardees, at all tiers. The Gonen laboratory is supported by funds from the Howard Hughes Medical Institute.

## Methods

### Materials

All commercial compounds were used as received without further recrystallization. **2**-**5** were purchased from InvivoChem LLC. **1** was purchased from Advanced ChemBlocks Inc.

### Grid preparation

Sample preparation followed the procedure as described previously.^24^ The carbon-coated copper grid (200-mesh, 3.05 mm O.D., Ted Pella Inc.) was pretreated with glow-discharge plasma at 15 mA on the negative mode for 45 s using PELCO easiGlow (Ted Pella Inc.). Less than 1 mg of each powdery compound were transferred and mixed with a grid in a 10 mL scintillation vial. After gently shaking the vial for 30 s, the grid was clipped using c-ring and autogrid clip (Thermo Fisher) at room temperature.

### MicroED data collection

The grids were loaded into the Thermo Fisher Talos Arctica TEM operating at 80 K and 200 keV (∼0.0251 Å wavelength). The TEM was pre-aligned using a 500 nm diffraction grating replica with latex spheres (Ted Pella Inc.). The CetaD camera (4096 × 4096 pixels) and EPUD software (Thermo Fisher) were used for MicroED data collection. The screening of microcrystals was done in the imaging mode (SA 5300x) (Fig. 2a-e). The MicroED datasets were collected in the diffraction mode at the calibrated sample to detector distance of 659 mm and the parallel beam condition (spot size 11, C2 aperture size 70 and C2 intensity 45.2%) which resulted in an illuminating dose rate of 0.01 e^-^/(Å^2^·s). The selected area aperture (resulting in ∼1.4 μm beam region) was inserted to cover the selected microcrystals during data collection to avoid diffraction from other samples in the beam. The typical data collection used a constant rotation rate of ∼2° per second over an angular wedge of 120° from -60° to +60°, with 0.5 s exposure time per frame. Prior to data collection, selected microcrystals were calibrated to eucentric heights to ensure that they remained inside the beam during the rotation.

### MicroED data processing

The MicroED movies were saved in mrc format and converted to smv format using the mrc2smv software (https://cryoem.ucla.edu/microed).^36^ The converted frames were indexed and integrated by XDS.^37^ The datasets with the highest resolution were scaled and merged using XSCALE,^37,38^ and their intensities were converted to SHELX hkl format using XDSCONV.^37,38^ The merged datasets showed increased completeness for **1**-**5** (from 83.2% to 99.8%), which could be *ab initio* solved by SHELXT^39^ or SHELXD^40^ (Supplementary Table 1). The structures were refined by SHELXL^41^ graphical interfae Shelxle^42^ to yield the final MicroED structures (Fig. 3 and Supplementary Table 1).

### Molecular Docking

The ligand structures of **1**-**3** and **5** were extracted from their MicroED structures and saved as mol2 files. Anions and water molecules were removed. Charges were kept based on the analysis of pKa values.^43^ Because **4** binds covalently with FAD coenzyme in MAO-A/B, therefore, **4**\* extracted from PDB entry 2VRM (resolution: 2.30 Å) was used as ligand.^44^ All these ligand structures were imported into AutoDock Tools 1.5.7,^45,46^ making all the active torsion bonds rotatable (Supplementary Table 2).

All the protein structures were downloaded from the Protein Data Bank (https://www.rcsb.org/). The structures of hSERT (PDB entry 5I73),^32^ MAO-A (PDB entry 2Z5X),^33^ and MAO-B (PDB entry 1OJ9)^34^ are determined by X-ray crystallography, while hNET (AF_AFP23975F1)^35^ and hDAT (AF_AFQ01959F1)^35^ are based on AlphaFold predictions with a global pLDDT score above 87. The solvent, ligand, and cofactor were removed using Pymol 2.5.5,^47^ except for the docking between **5** and MAO-A/B where the FAD was kept in the original position. After that, hydrogen atoms and charges were computed and added via AutoDock Tools 1.5.7 (Supplementary Table 2).^45,46^

Template structures were obtained in PDB database via the query function in CB-Dock2 webtool^48^ or literatures, considering the similarities in protein and ligand (Supplementary Fig. 11). The approximate docking center was determined and positioned with an 18.75 Å × 18.75 Å × 18.75 Å grid box in AutoDock Vina1.1.2.^45,46^ During the molecular docking, all active torsion bonds in the ligands were set to be flexible, and the receptor was set to be rigid. The docked complex with the lowest binding energy was analyzed by the Protein-Ligand Interaction Profiler (PLIP) web tool.^49^

## Notes

### Competing Interest Statement

The authors have declared no competing interest.

