## Supplementary Information for "Uncovering the Elusive Structures and Mechanisms of Prevalent Antidepressants"

**Supplementary Table 1** MicroED data statistics of five antidepressants.[a]

| Compound | **1** | **2** | **3** | **4** | **5** |
|---|---|---|---|---|---|
| Name | Reboxetine mesylate | Pipofezine dihydrochloride monohydrate | Ansofaxine hydrochloride dihydrate | Phenelzine sulfate | Bifemelane hydrochloride |
| Stoichiometric formula | $C_{20}H_{27}NO_6S$ | $C_{16}H_{21}Cl_2N_5O_2$ | $C_{24}H_{36}ClNO_5$ | $C_8H_{14}N_2O_4S$ | $C_{18}H_{24}ClNO$ |
| Mr | 409.50 | 388.30 | 454.03 | 234.28 | 305.80 |
| Temperature (K) | 80 | 80 | 80 | 80 | 80 |
| Crystal system | Monoclinic | Monoclinic | Monoclinic | Monoclinic | Orthorhombic |
| Space group | P $2_1$/c | P $2_1$/c | P $2_1$/c | P $2_1$/c | P bca |
| Unit cell lengths (Å) | | | | | |
| a | 20.000(4) | 6.880(2) | 14.800(3) | 20.360(4) | 12.870(3) |
| b | 5.490(2) | 15.610(3) | 10.270(2) | 5.460(2) | 7.310(3) |
| c | 19.060(4) | 15.930(3) | 16.040(3) | 20.300(4) | 35.100(7) |
| Unit cell angles (°) | | | | | |
| $\alpha$ | 90.00(3) | 90.00(3) | 90.00(3) | 90.00(3) | 90.00(3) |
| $\beta$ | 107.34(3) | 97.22(3) | 95.31(3) | 111.21(3) | 90.00(3) |
| $\gamma$ | 90.00(3) | 90.00(3) | 90.00(3) | 90.00(3) | 90.00(3) |
| Cell volume (Å$^3$) | 1997.7(10) | 1697.3(7) | 2427.6(8) | 2103.8(10) | 3302.2(14) |
| No. of datasets merged | 7 | 4 | 3 | 1 | 3 |
| No. of observed reflections | 93185 | 32120 | 33547 | 21752 | 39392 |
| No. of unique reflections | 5384 | 3356 | 4068 | 7358 | 3423 |
| $R_{obs}$ (%) | 27.7 | 33.6 | 28.8 | 17.2 | 28.2 |
| $R_{meas}$ (%) | 28.6 | 35.5 | 30.8 | 21.2 | 29.6 |
| I/Sigma | 8.13 | 5.20 | 5.22 | 3.99 | 5.78 |
| $CC_{1/2}$ | 99.4 | 97.8 | 98.1 | 98.9 | 98.9 |
| Completeness (%) | 99.8 | 99.7 | 86.3 | 83.2 | 97.7 |
| **Resolution (Å)** | **0.73** | **0.82** | **0.83** | **0.63** | **0.83** |
| **$R_1$ (%)** | **19.3** | **15.9** | **18.7** | **20.2** | **19.6** |
| $wR_2$ (%) | 52.1 | 40.2 | 41.1 | 48.7 | 47.1 |
| GooF | 2.134 | 1.439 | 1.422 | 1.423 | 1.712 |

**Notes:** MicroED structures **1** and **3-5** were solved by SHELXT,[1] structure **2** was solved by SHELXD;[2] All MicroED structures were refined by SHELXL.[3]

**Supplementary Table 2** Molecular Docking of five antidepressants and their target proteins

| Ligand[a] | Protein[b] | Cofactor/ Coenzyme | Template PDB ID[5-10] | Binding site[c] |
|---|---|---|---|---|
| **1R** | hNET | No | 4XNX | S1 |
| **1R** | hNET | No | 3GWV | S2 |
| **1S** | hNET | No | 4XNX | S1 |
| **1S** | hNET | No | 3GWV | S2 |
| **2** | hSERT | No | 5I73 | near S2 |
| **3R** | hSERT | No | 5I73 | S1 |
| **3R** | hNET | No | 4XNX | S1 |
| **3R** | hDAT | No | 4MMC | S1 |
| **3S** | hSERT | No | 5I73 | S1 |
| **3S** | hNET | No | 4XNX | S1 |
| **3S** | hDAT | No | 4MMC | S1 |
| **4*** | MAO-A | No | 2Z5X | C1 |
| **5** | MAO-A | Yes | 2Z5X | R1 |
| **5** | MAO-B | Yes | 1OJ9 | R1′ |

Notes: [a]The ligand structures were directly extracted from MicroED structures of **1-3** and **5**, while the ligand structure of **4*** was extracted from PDB structure 2VRM (resolution: 2.30 Å). [b]The protein structures were obtained from PDB database either by X-ray crystallography or AlphaFold, hNET (AF_AFP23975F1),[4] hSERT (5I73),[7] hDAT (AF_AFQ01959F1),[4] MAO-A (2Z5X),[9] MAO-B (1OJ9).[10] [c]The binding sites (docking center) was determined by aligning the protein with template structures, which was then used within an 18.75 Å × 18.75 Å × 18.75 Å grid box in AutoDock Vina 1.1.2.[11,12]

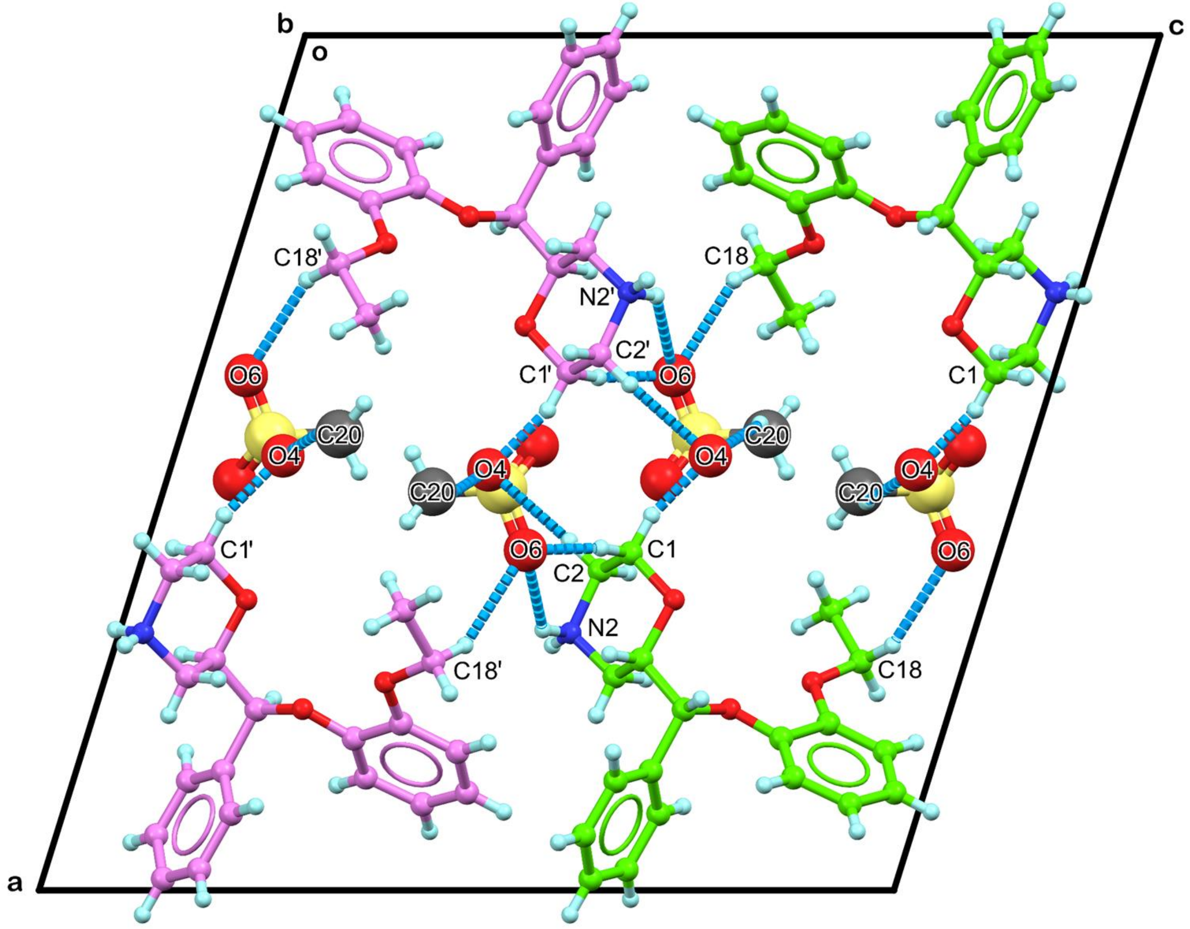


**Supplementary Figure 1** The packing diagram of **1**, viewed along the *b*-axis. Hydrogen bonding and selected dipole-dipole interactions were represented by the dashed lines in marine, **1R** was colored in green, **1S** was colored in violet.

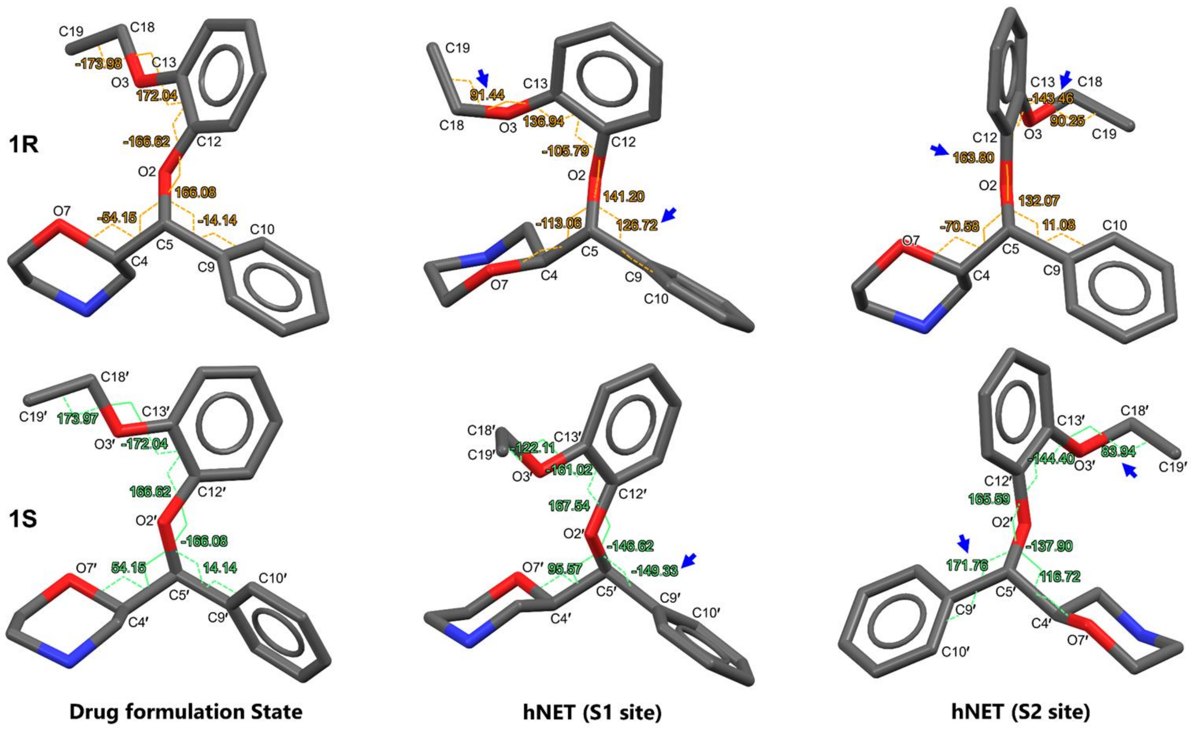


**Supplementary Figure 2** The major conformational changes between the crystal structures of **1** and their molecular docking structures in hNET. The major torsion changes were highlighted with blue arrows.

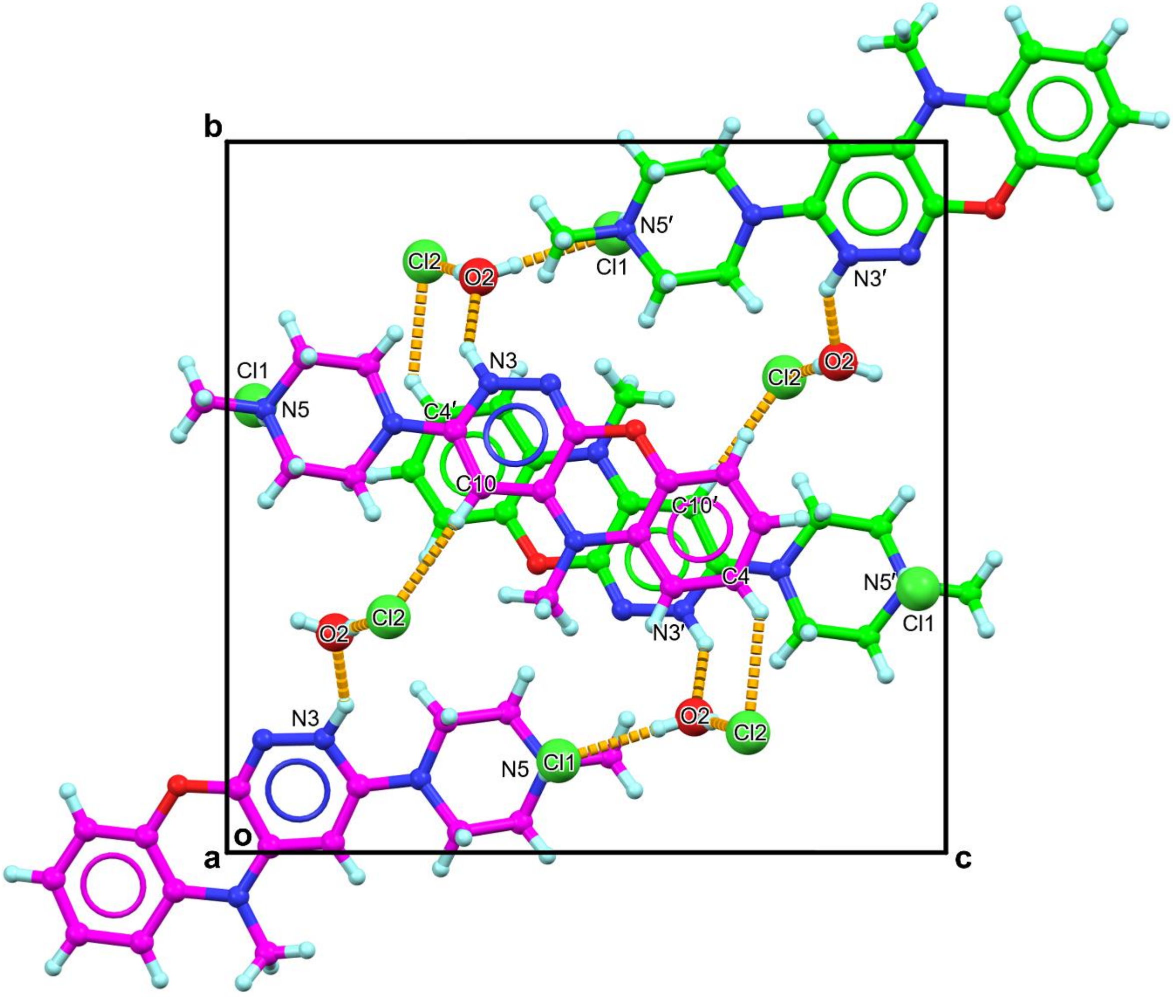


**Supplementary Figure 3** The packing diagram of **2**. Hydrogen bonding and selected dipole-dipole interactions were represented by the dashed lines in orange. **2a** was colored in magenta, **2b** was colored in green.

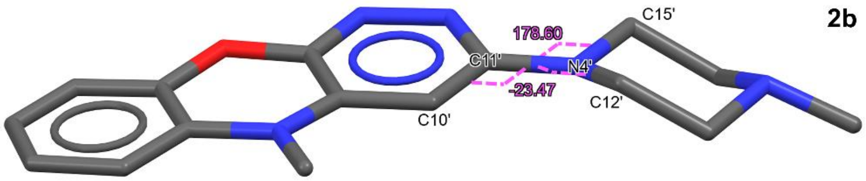


**Drug formulation State**

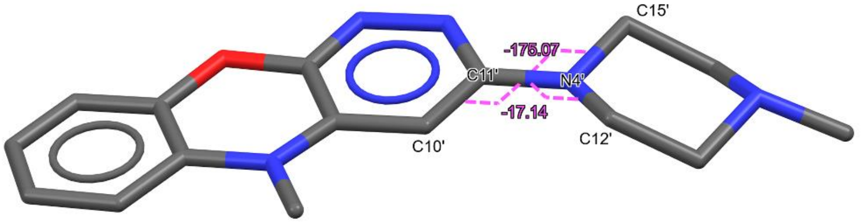


**hSERT (near S2 site)**

**Supplementary Figure 4** The conformational changes between the crystal structure of **2b** and its molecular docking structure in hSERT. The freely rotatable torsion angles (C10′–C11′–N4′–C12′ and C10′–C11′–N4′–C15′) were measured.

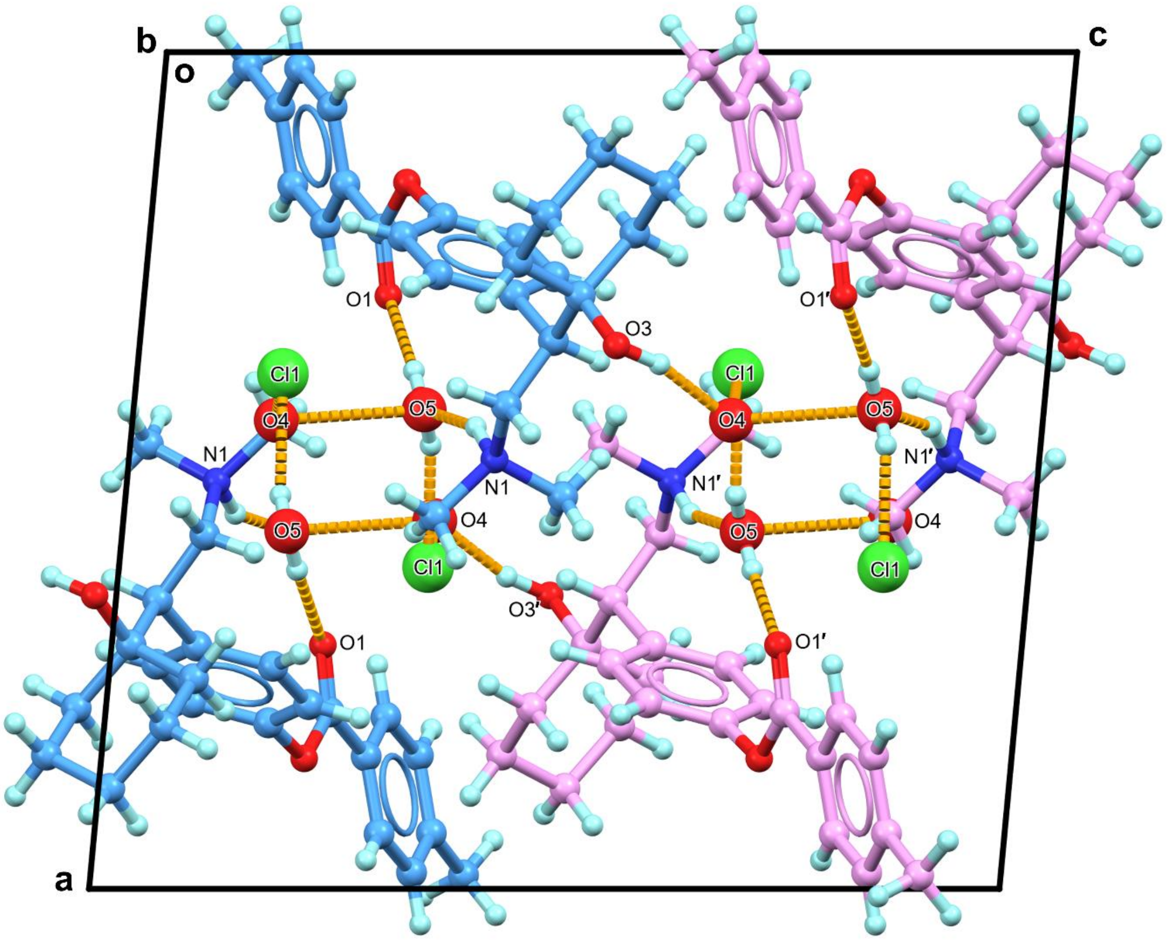


**Supplementary Figure 5** The packing diagram of **3**, viewed along the *b*-axis. Hydrogen bonding interactions were represented by the dashed lines in orange, **3R** was colored in blue, **3S** was colored in violet. The Cl1–O4–O5 is measured at 107.7° and O1–O5–Cl1 is measured at 112.8°.

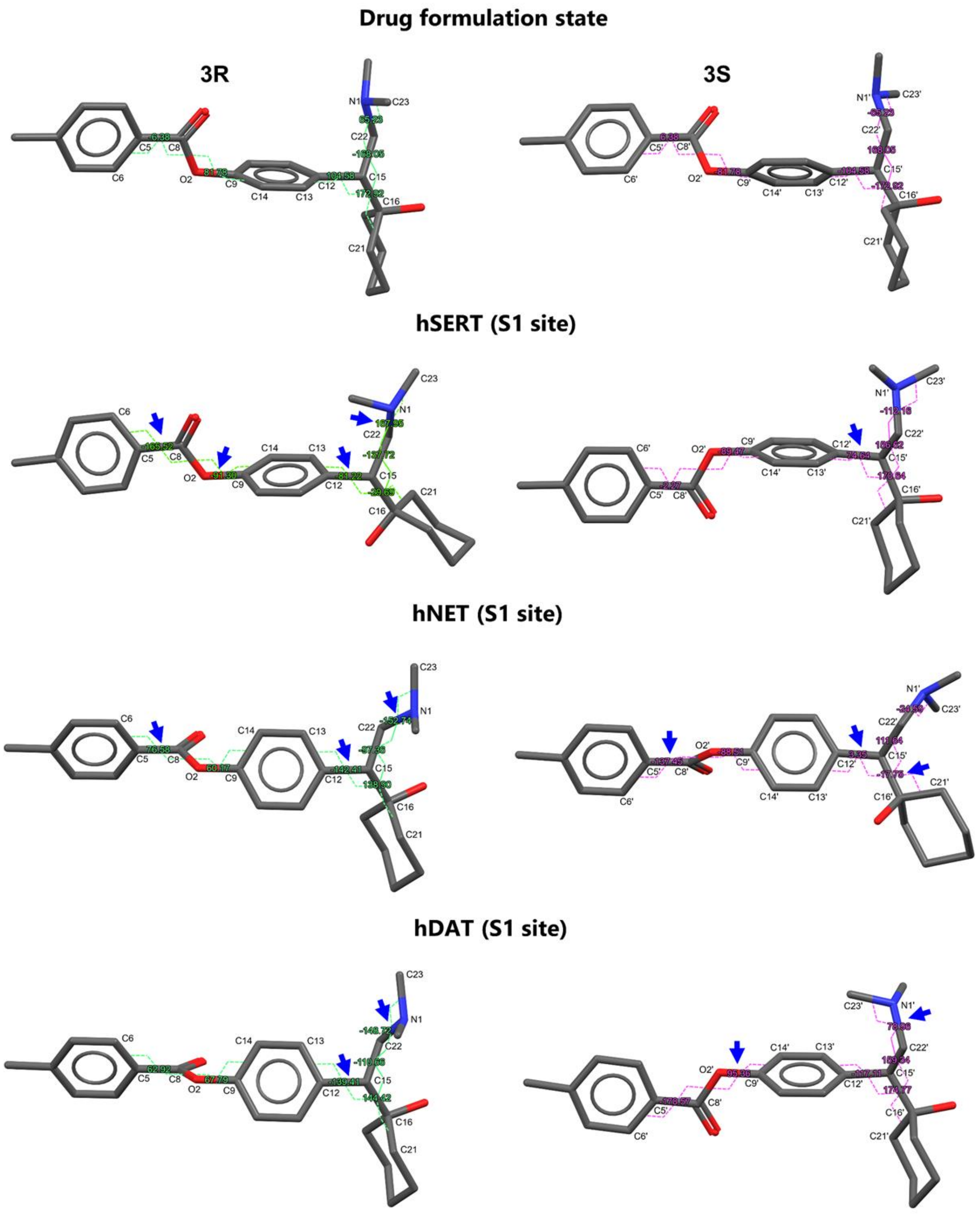


**Supplementary Figure 6** The major conformational changes between the crystal structures of **3** and their molecular docking structures in hSERT, hNET and hDAT. The major torsion changes were highlighted with blue arrows.

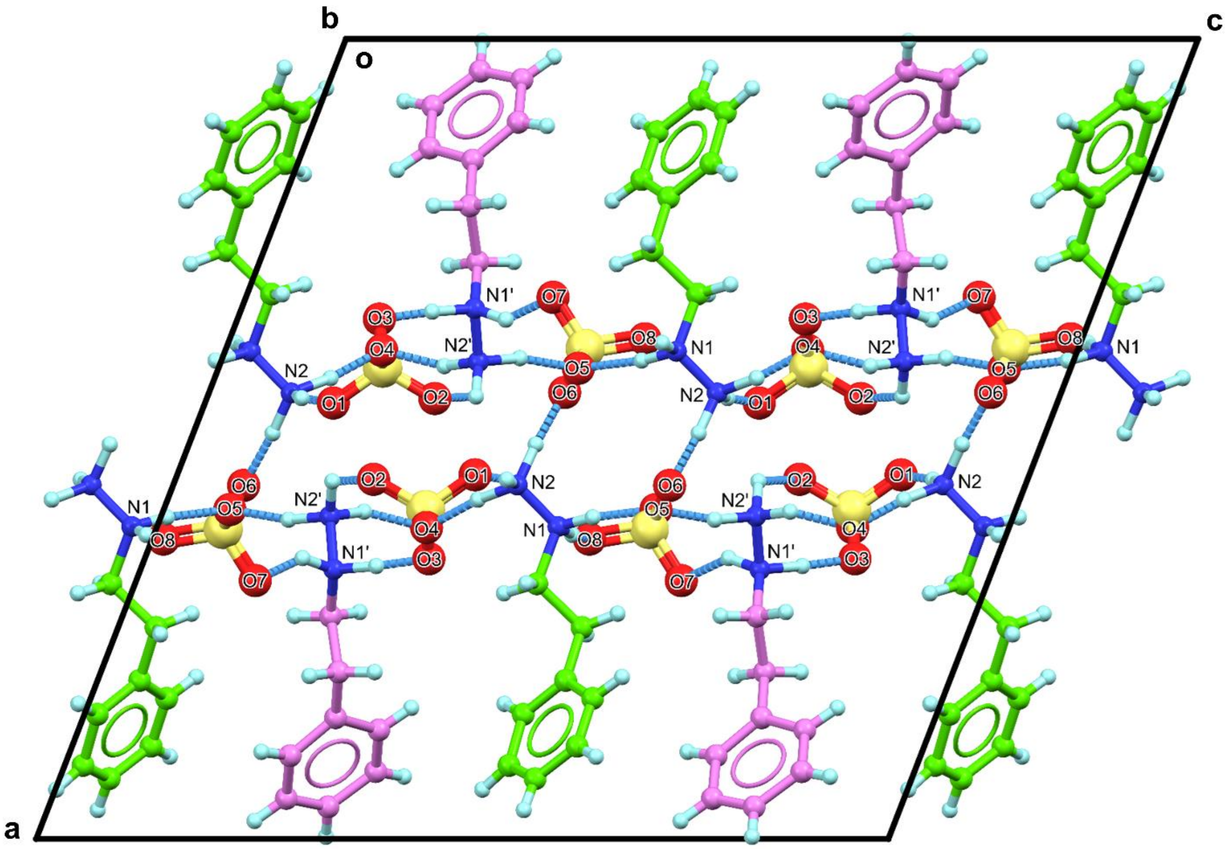


**Supplementary Figure 7** The packing diagram of **4**, viewed along the *b*-axis. **4a** was colored in green, **4b** was colored in violet. Hydrogen bonding interactions were represented by the dashed lines in marine.

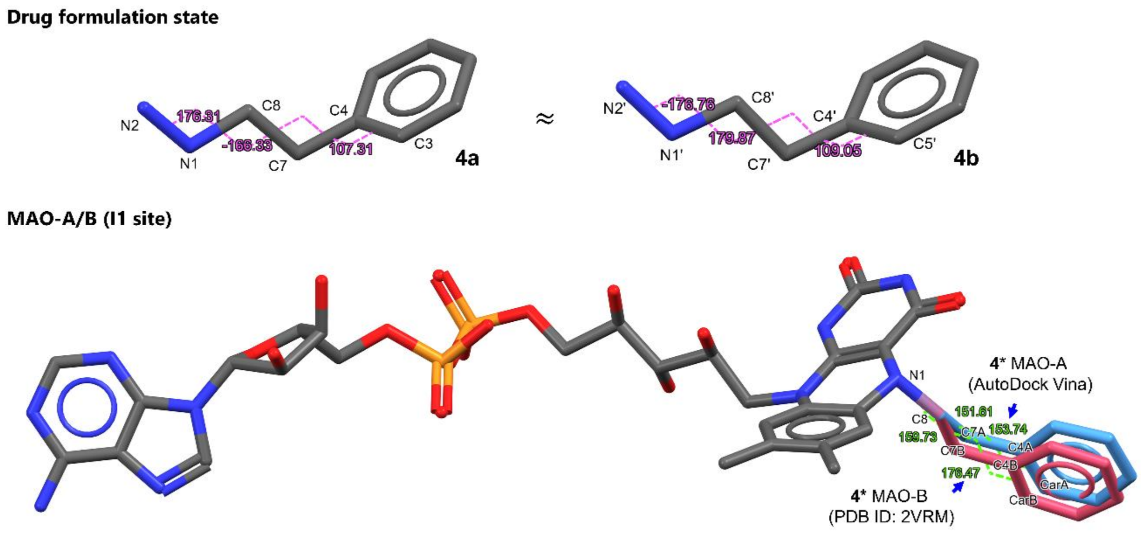


**Supplementary Figure 8** The major conformational changes between the crystal structures of **4** and their docking structures **4*** in MAO-A/B. Phenelzine covalently linked to the FAD in C1 site of MAO-A/B as **4***. Selected torsion angles (C3–C4–C7–C8, C4–C7–C8–N1, C7–C8–N1–N2 in **4a**, C5′–C4′–C7′–C8′, C4′–C7′–C8′–N1′, C7′–C8′–N1′–N2′ in **4b**, $C_{ar}$A–C4A–C7A–C8, C4A–C7A–C8–N1 and $C_{ar}$B–C4B–C7B–C8, C4B–C7B–C8–N1 in **4*** in MAO-A/B) were measured. In the biological state, only the phenylethyl part will covalently bind to the FAD cofactor in MAO-A/B, where the C4–C7/C4′–C7′ is presumed to have 44° to 69° rotations. The major torsion changes were highlighted with blue arrows.

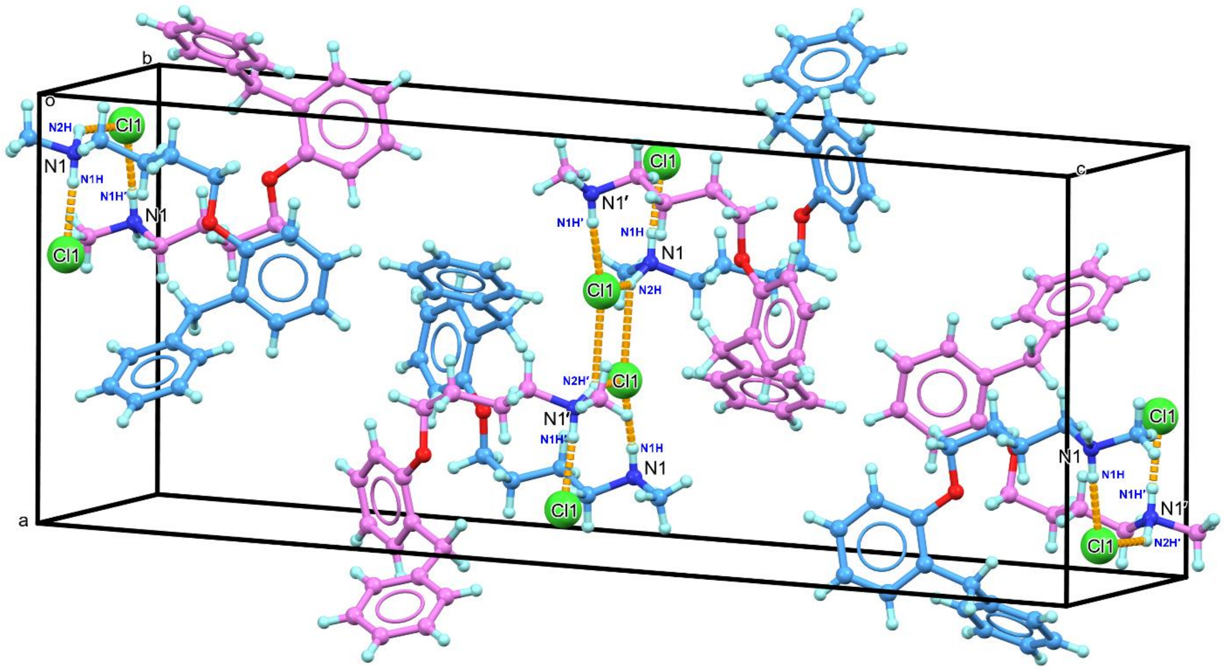


**Supplementary Figure 9** The packing diagram of **5**, viewed along the *b*-axis. **5a** was colored in blue, **5b** was colored in violet. Hydrogen bonding interactions were represented by the dashed lines in orange.

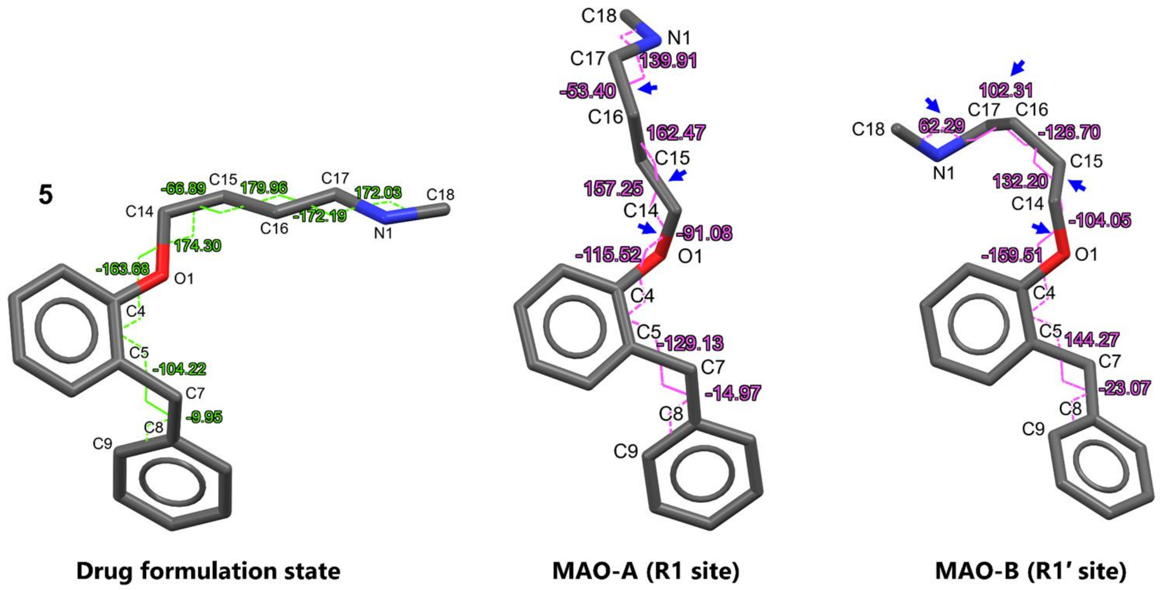


**Supplementary Figure 10** The major conformational changes between the crystal structures of **5** and its molecular docking structures in MAO-A/B. The major torsion changes were highlighted with blue arrows.

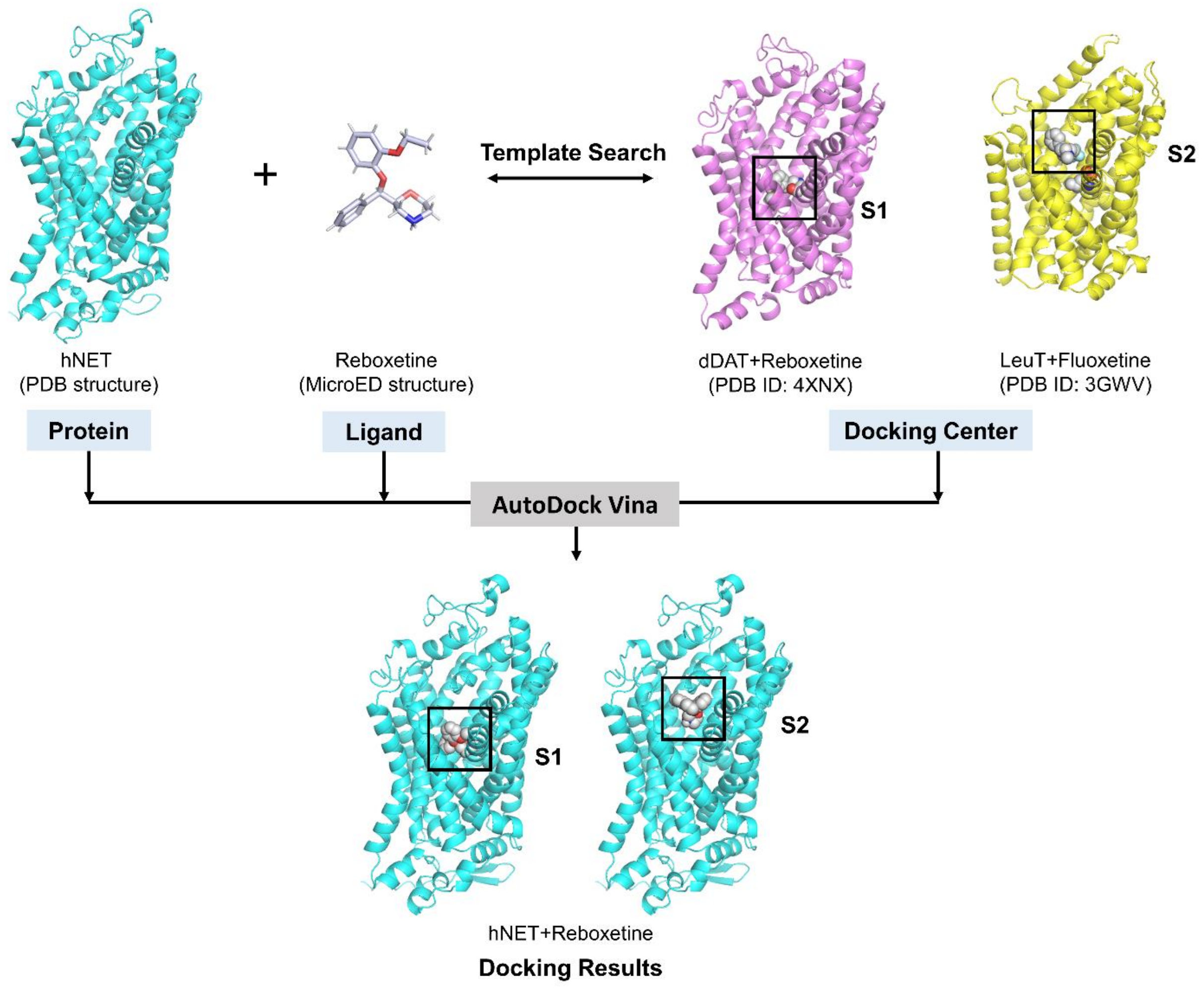


**Supplementary Figure 11** AutoDock Vina docking workflow illustrated by hNET and Reboxetine. The template structures were obtained in PDB database via the query function in CB-Dock2 webtool[13] or searching the literatures, considering the similarities of both protein and ligand. The docking center was positioned by aligning the protein with template structures, and then used within an 18.75 Å × 18.75 Å × 18.75 Å grid box in AutoDock Vina 1.1.2.[11,12]

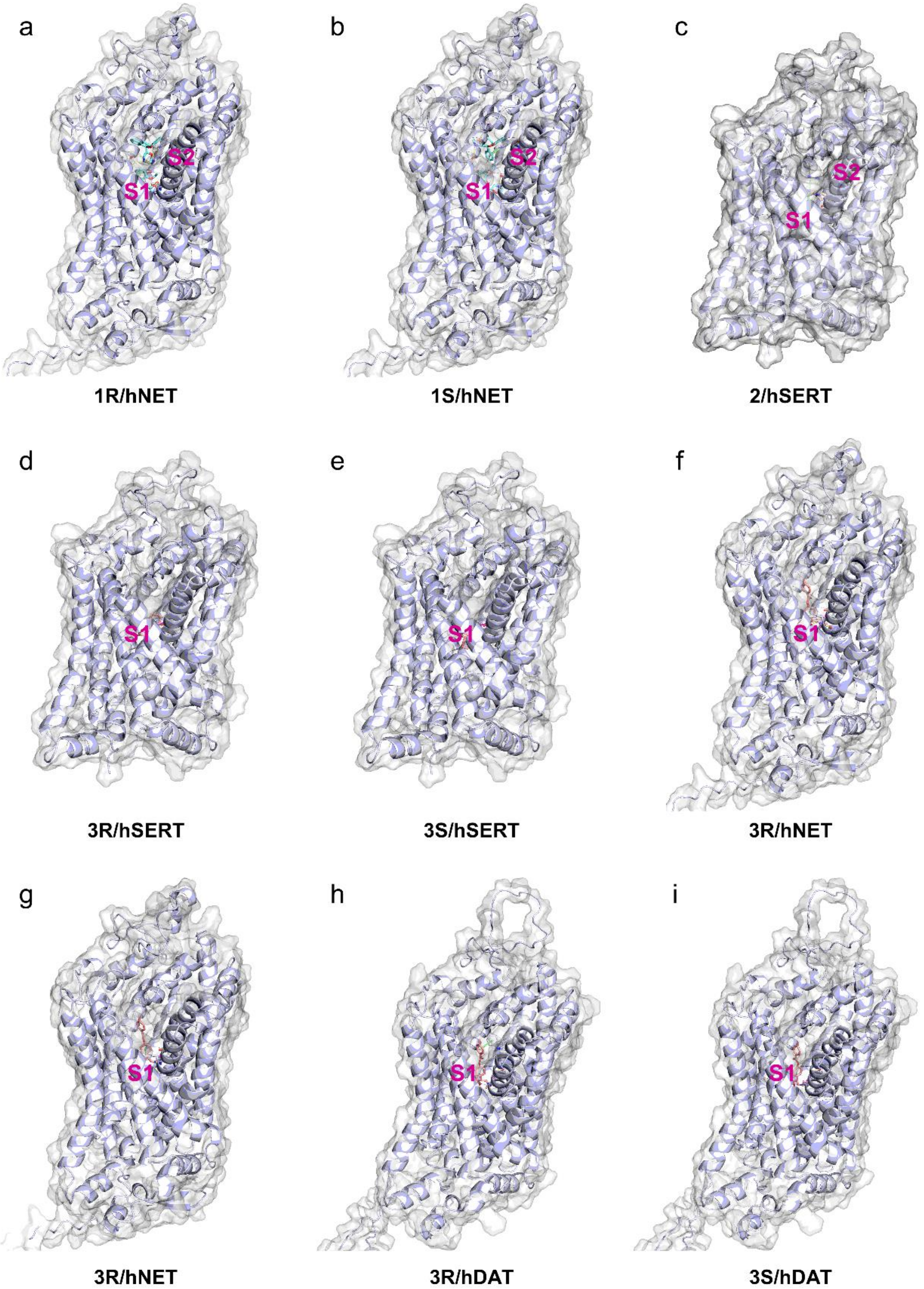


**Supplementary Figure 12** Overall view of protein-drug interaction diagram of complex between (a) **1R** and hNET; (b) **1S** and hNET; (c) **2** and hSERT; (d) **3R** and hSERT; (e) **3S** and hSERT; (f) **3R** and hNET; (g) **3S** and hNET; (h) **3R** and hDAT; (i) **3S** and hDAT. The binding sites were marked.

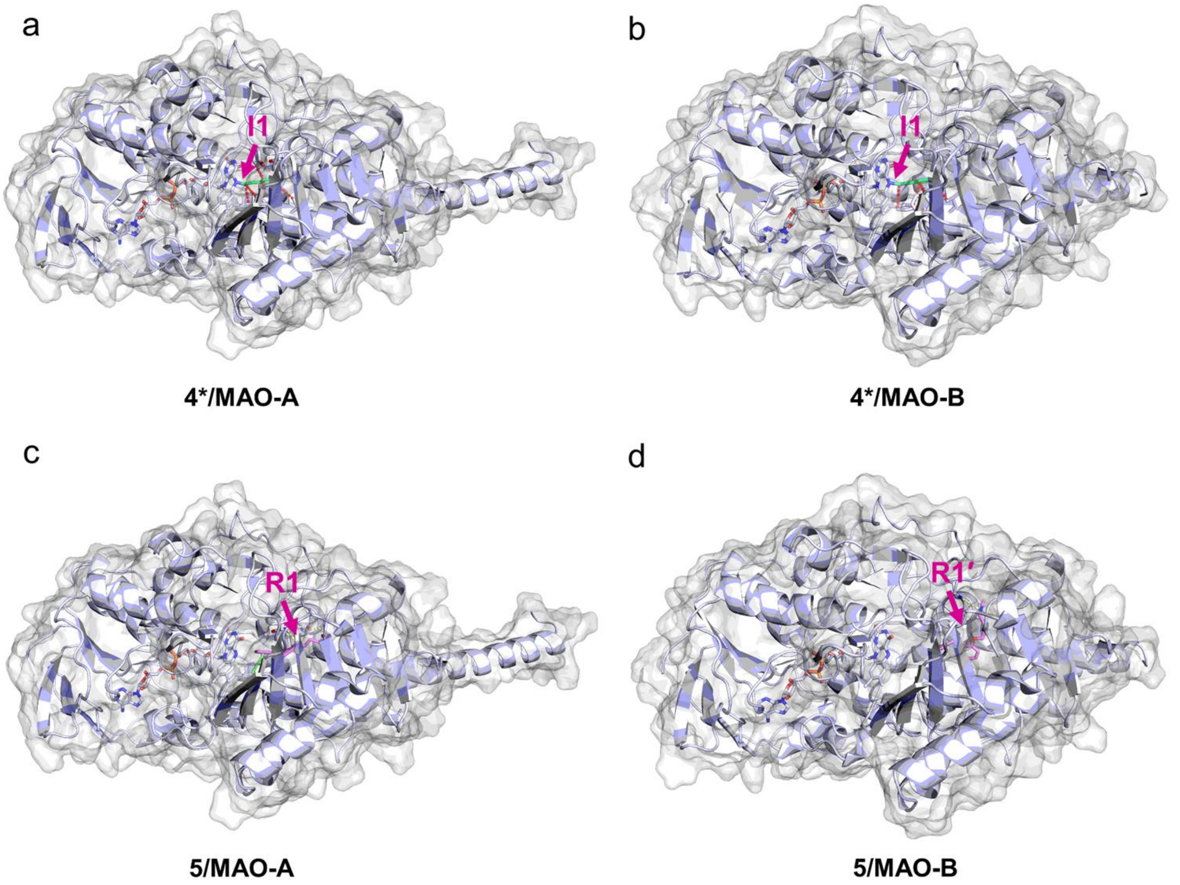


**Supplementary Figure 13** Overall view of protein-drug interaction diagram of complex between (a) **4*** and MAO-A; (b) **4*** and MAO-B; (c) **5** and MAO-A; (d) **5** and MAO-B. The binding sites were marked.

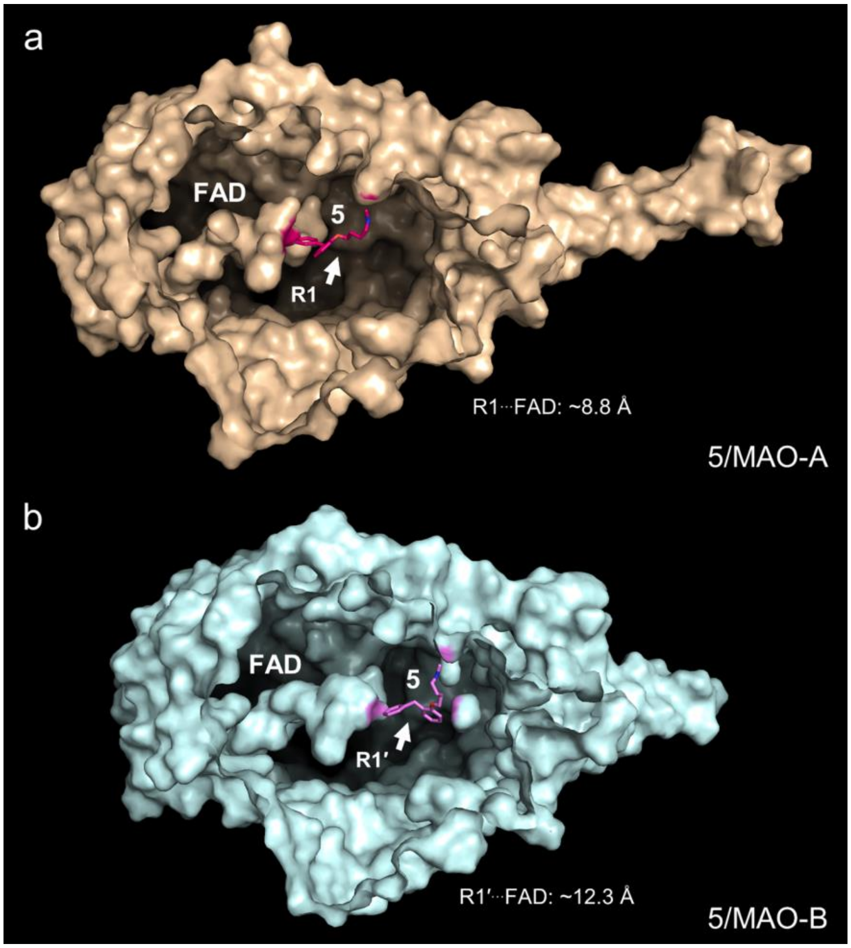


**Supplementary Figure 14** Protein voids observed in **5**/MAO-A and **5**/MAO-B complexes. The distances between FAD and **5** were measured and compared.
